# The structure of hippocampal circuitry relates to rapid category learning in humans

**DOI:** 10.1101/2021.05.14.444187

**Authors:** Margaret L. Schlichting, Melisa Gumus, Teresa Zhu, Michael L. Mack

**Author notes:** **Corresponding Author:** Michael L. Mack, Department of Psychology, University of Toronto, 100 St. George Street, 4. Authors made equal contributions (Margaret L. Schlichting, Michael L. Mack). **Data availability:** Upon publication, data related to this publication will be available on OSF - https://osf.io/vwyqg/?view_only=5259728d722d4a38a5b73e238bf1f385.

## Abstract

Prior work suggests that complementary white matter pathways within the hippocampus differentially support learning of specific versus general information. In particular, while the trisynaptic pathway (TSP) rapidly forms memories for specific experiences, the monosynaptic pathway (MSP) slowly learns generalities. However, despite the theorized significance of such circuitry, characterizing how information flows within hippocampus (HPC) to support learning in humans remains a challenge. We leveraged diffusion-weighted imaging as a proxy for individual differences in white matter structure linking key regions along TSP (HPC subfields CA_3_ and CA_1_) and MSP (entorhinal cortex and CA_1_) and related these differences in hippocampal structure to category learning ability. We hypothesized that learning to categorize the “exception” items that deviated from category rules would benefit from TSP-supported mnemonic specificity. Participant-level estimates of TSP- and MSP-related integrity were constructed from HPC subfield connectomes of white matter streamline density. Consistent with theories of TSP-supported learning mechanisms, we found a specific association between the integrity of CA_3_-CA_1_ white matter connections and exception learning. These results highlight the significant role of HPC circuitry in complex human learning.

## Introduction

Our memories must contain both specifics of individual events and generalizations that span experiences to be maximally useful. Longstanding theories (McClelland et al., 1995) on the complementary nature of hippocampal and cortical memory processes have highlighted the role of the hippocampus—a structure long known to be critical for normal episodic memory (Scoville and Milner, 1957)—as one of recording the specific details of individual episodes. However, recent modelling work suggests that the hippocampus itself is capable of forming both specific and general memories (Schapiro et al., 2017)—but that critically, it may do so through distinct anatomical pathways that exist *within* the structure (Norman and O’Reilly, 2003; Ketz et al., 2013).

The monosynaptic (MSP) and trisynaptic (TSP) intra-hippocampal pathways are thought to underlie the relatively slower (Gall et al., 1998; Lee and Kesner, 2002; Rolls and Kesner, 2006) accumulation of generalizations and rapid storage of specifics, respectively (Lee et al., 2004; Nakashiba et al., 2008). Such functional differences might be the product of the distinct anatomy of these pathways: MSP connects inputs from entorhinal cortex (ERC) to the *cornu ammonis* 1 (CA_1_) subfield of hippocampus directly, while such connections in TSP are via dentate gyrus and CA_3_. The inclusion of DG and CA_3_ in the TSP intra-hippocampal pathway is thought to underlie the more precise, relational representations that facilitate pattern completion-based learning. Therefore, it might be the case that the structural integrity of one’s TSP—rather than other features like hippocampal volume—might be most predictive of learning, particularly when such learning primarily taxes TSP. However, to our knowledge the integrity of connections between subfields along the TSP has been neither measured nor related to behaviour among healthy young adults using diffusion weighted imaging (DWI), leaving a large gap between theoretical frameworks highlighting the importance of intra-hippocampal pathways and empirical studies largely focussing on hippocampal volume (Chadwick et al., 2014; Travis et al., 2014; Daugherty et al., 2016; Canada et al., 2019) or morphology (Voineskos et al., 2015; DeKraker et al., 2020).

Recent work has begun to appreciate the widespread role of the hippocampus in cognition (Shohamy and Turk-Browne, 2013), including in the domain of category learning (Mack et al., 2018; Zeithamova et al., 2019). Building upon previous research showing the involvement of hippocampus (Davis et al., 2012) and its representations (Mack et al., 2016; Bowman and Zeithamova, 2018) in supporting such behaviours, here we asked whether performance variability across individuals (Little and McDaniel, 2014; Shen and Palmeri, 2016) might be explained by the integrity of connections between TSP-related regions in particular. We set up our category structure such that while a simple similarity-based calculation would yield successful categorization for *most* items, some items—termed “exceptions”—violate the simple rule and therefore must be learned separately. Informed by the predictions of a computational model (SUSTAIN; Love et al., 2004; Love and Gureckis, 2007) that we and others have previously shown is reflected in neural response (Davis et al., 2012; Mack et al., 2016, 2020), we reasoned that exceptions might preferentially be supported by TSP-based mechanisms. In particular, SUSTAIN suggests that exception learning is accomplished by computing the discrepancy (i.e., mismatch) between the stored memories that are reactivated—that is, pattern completed, presumably through CA_3_-based operations (Neunuebel and Knierim, 2014)—and the current experience in light of corrective feedback (Sakamoto and Love, 2004; Davis et al., 2012).

Because exceptions are by design more similar to the *alternate* category, such pattern completion will yield retrieval of alternate-category memories and incorrect categorization. It is this mismatch computed from the feedback that will therefore promote encoding of these exception items.

In the context of this theoretical framework, we set out to test the hypothesis that the ability to learn to successfully categorize exception items—but not rule-following ones—would be accounted for in part by the structural integrity of TSP-related—but not MSP-related— connections as measured with DWI. We leveraged sophisticated automated tools both for segmenting HPC subfields and medial temporal lobe (MTL) cortex (Yushkevich et al., 2015) and for estimating structural connectivity (Smith et al., 2012; Tournier et al., 2019) between key regions along TSP (CA_3_ to CA_1_) and MSP (ERC to CA_1_). This approach potentially sacrifices precision relative to more traditional manual methods for segmentation and tractography, but provides a more efficient and flexible data-driven approach for investigating brain-behaviour relationships in larger samples. Importantly, in employing these methods, we find that the density of TSP-related connections (CA_3_ to CA_1_) are associated with individual variability in exception learning.

## Results

Thirty-seven healthy young adults performed a feedback-based category learning task in which they classified a set of highly similar visual stimuli (flowers) into two categories (shade- or sun-preferring). Stimuli varied along three dimensions (inner petal shape, outer petal shape, and petal colour; **Fig. 1A**), but the category structure was such that there was no simple rule that separated the categories. Rather, each category included an “anchor” item, two “similar” items (which differed from the anchor on one dimension), and one “exception” item (which differed on two dimensions) (Shepard, 1961; Nosofsky et al., 1994). Critically, while categorization of anchor and similar items could be based on visual features alone, exception items were in fact more similar to the anchor for the *alternate* category (**Fig. 1B**). We hypothesized that because participants would need to form separate, specific memories for how to classify these non-rule-following items, rapid encoding mechanisms would be uniquely related to categorization task (**Fig. 1C**) performance for exceptions.

**Figure 1:**
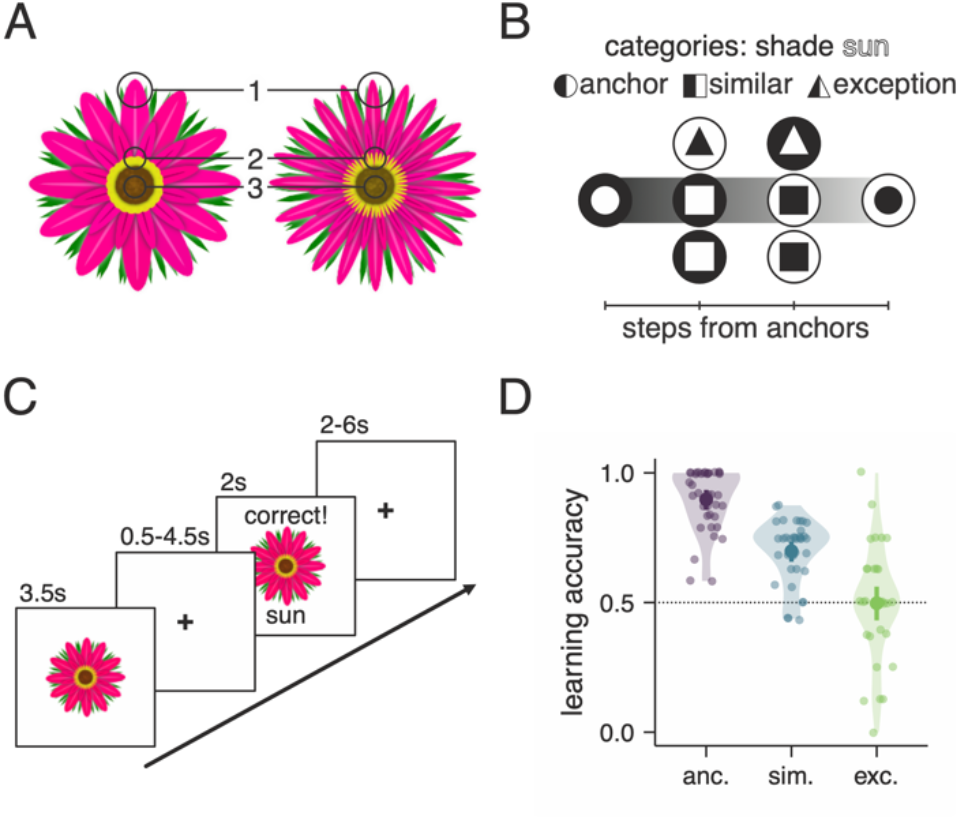
Example stimuli, task schematic, and learning performance. **A**) Flower stimuli varied in (1) width of outer (pink) petals, (2) shape of inner (yellow) petals, and (3) colour and texture of brown centre. **B**) Subway plot depicting all task stimuli as a function of distance from both shade- and sun-preferring flower anchors (circles at left- and rightmost positions on map, respectively). Shade-preferring flowers are denoted with dark outer circle; sun-preferring denoted with white outer circle. Similar items (squares) are one step in feature space (i.e., one “subway stop”) away from their own category anchor. Exception items (triangles) are two steps away from their own category anchor, and only one step from the *opposite* category anchor. **C**) Participants decided whether an individual flower presented on the screen was either sun- or shade-preferring, and then received feedback about their choice. **D**) Final categorization performance, operationalized as the average accuracy in categorizing anchor (anc.), similar (sim.), and exception (exc.) items during the last third of the category learning task.

End of learning performance (**Fig. 1D**) differed by item type (χ^2^_2_=194.07, p=7.224×10^-43^), being highest (and near-perfect) for anchors, lowest for exceptions, and intermediate for similar items, as expected (Davis et al., 2012). With evidence of all participants showing learning for the more frequently presented anchors, we focus the following analyses on similar and exception items which were matched to one another in terms of number of presentations. Importantly, participants exhibited a range of performance for both similar (range: 43.75-87.5%) and exception (0-100%) items, allowing us to ask how individual differences in brain structure— particularly, integrity of the connections among hippocampal subfields—relates to behaviour.

Using automatically defined (Yushkevich et al., 2015) hippocampal and MTL cortical substructures (Hindy et al., 2016) in combination with DWI, we generated tractography-based estimates of the integrity of white matter connections associated with MSP (operationalized as streamline count from ERC-CA_1_) and TSP (CA_3_-CA_1_) for each participant (**Fig. 2**). Our reasons for defining the pathways in this way—including only part of the overall TSP pathway (i.e., excluding ERC-DG, DG-CA_3_, and output connections via the fornix)—were to (1) align with prior computational work which has focussed on the CA_3_-CA_1_ connection as reflecting TSP strength (Ketz et al., 2013; Schapiro et al., 2017) and (2) ensure that the two pathways were defined with a similar number of constraints such that neither pathway was at a clear disadvantage (Baum et al., 2018). We additionally extracted in-scanner motion summary estimates for each DWI scan to control for the possibility that person-to-person variability in the ability to remain still for the duration of the scan could impact our estimates of path integrity. All reported analyses include motion as a covariate, but results were virtually identical when motion was ignored. In addition, while our main statistical analyses require scaling of the streamline counts, we display the raw streamline counts by pathway in **Fig. 2B**.

**Figure 2:**
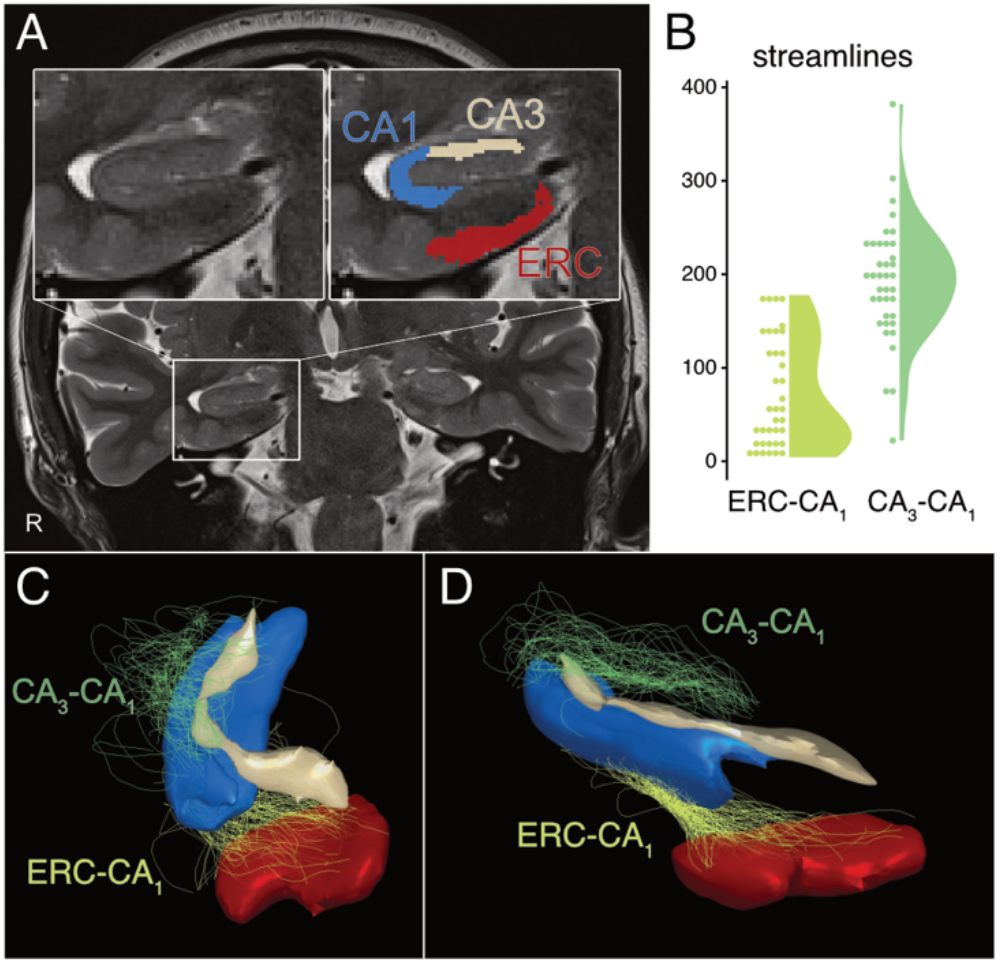
Example segmentation and tractography from one sample participant. **A**) Coronal slice of the brain focussed on the participant’s MTL. Insets show close-up view of hippocampus (left) with superimposed ROIs (right) used as masks in the present tractography analysis (blue, CA_1_; red, ERC; cream, CA_3_). **B**) Distribution of raw streamline counts across all participants, shown as both dot plots (left, where each dot represents a single participant), and violins (right, representing the smoothed distribution). These values were scaled for subsequent statistical analyses to yield the values plotted along the x-axis in **Fig. 3A**. **C**) and **D**) show two different views of CA_3_-CA_1_ and ERC-CA_1_ streamline renderings in 3D to enable visualization of the pathways for the participant shown in panel **A**. Regions of interest (smoothed) are coloured as in panel A, and other structures have been removed for easy visualization. In panels **B-D**, light green denotes ERC-CA_1_ and dark green denotes CA_3_-CA_1_. See supplement **Fig. S1** for additional depictions of streamline estimates.

When both tracts were considered simultaneously in a multiple regression, tract integrity of CA_3_-CA_1_ (regression coefficient β=0.117, CI_95%_=[0.031, 0.203], p=0.010) but not ERC-CA_1_ (β=− 0.038, CI_95%_=[−0.144, 0.068], p=0.472) was related to individual differences in performance for the exception items (**Fig. 3**)^1^. The reader will note that because our CA_3_-CA_1_ and ERC-CA_1_ values were scaled, the regression coefficients can be interpreted as the increase in behaviour associated with a one standard deviation (SD) increase in streamline count. In other words, exception learning performance increases by about 10% for every SD increase in CA_3_-CA_1_ integrity. The reliability of the relationship between exception learning and CA_3_-CA_1_ integrity was confirmed with bootstrap resampling (**Fig. 3B;** p_boot_=0.010) and robust regression analyses (p=0.004). The interaction was also significant (β=0.180, CI_95%_=[0.044, 0.317], p=0.010), such that the relationship between accuracy and tract integrity was significantly stronger for CA_3_-CA_1_ than ERC-CA_1_. Importantly, this brain-behaviour relationship was only observed for exception items; neither CA_3_-CA_1_ nor ERC-CA_1_ tract integrity was associated with performance on similar items (both p>0.35).

**Figure 3:**
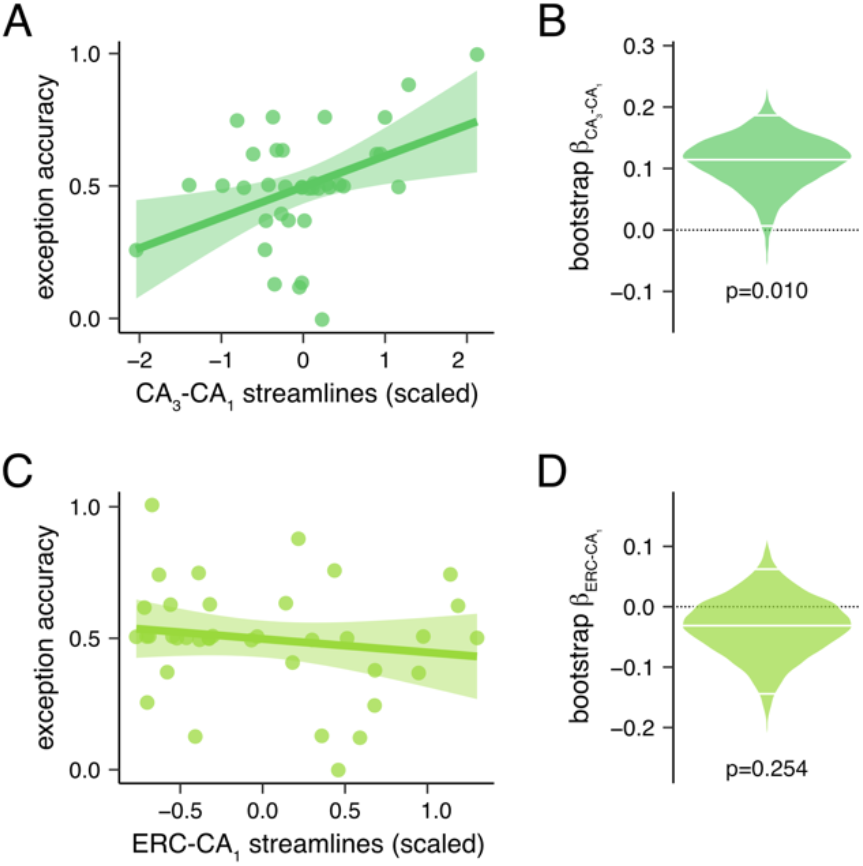
Individual differences in CA_3_-CA_1_ but not ERC-CA_1_ tract integrity are related to exception performance. **A**) Simple linear relationships between tract integrity (x-axes) of CA_3_-CA_1_ (top, dark green) and ERC-CA_1_ (bottom, light green) and final categorization performance for exception items (y-axes). CA_3_-CA_1_ (regression coefficient β=0.122, CI_95%_=[0.037, 0.206], p=0.006) but not ERC-CA_1_ (β=-0.061, CI_95%_=[−0.175, 0.053], p=0.284) tract integrity shows a significant relationship with performance. Note that while results in the main paper focus on a multiple regression in which the relationship between CA_3_-CA_1_ and performance remains when controlling for ERC-CA_1_, here we depict the simple linear regressions for visualization purposes. **B**) Distributions of bootstrapped regression coefficients; white lines depict median and 95% confidence intervals of β values, in which participants were randomly sampled with replacement across 10,000 iterations. The CA_3_-CA_1_ effect was reliable across bootstrapped samples (CI_95%_=[0.024,0.187], p_boot_=0.010), the ERC-CA_1_ effect was not (CI_95%_=[−0.149,0.065], p_boot_=0.254).

To assess the specificity of this relationship to intra-hippocampal path integrity as opposed to more gross anatomical features like structure size, we additionally tested for possible relationships between MTL volumes and performance. There was no relationship between the volume of any MTL substructure (CA_1_, CA_3_, DG, SUB, ERC) and performance for either exception (all p>0.54) or similar (all p>0.16) items when only volume was considered. Importantly, the relationship between CA_3_-CA_1_ tract integrity and exception item performance remained reliable (β=0.129, CI_95%_=[0.021, 0.237], p=0.021) when controlling for individual differences in volume of hippocampal subfields (CA_1_, CA_3_, DG, SUB) and entorhinal cortex (ERC); none of the other relationships were significant (ERC-CA_1_: β=-0.034, CI_95%_=[−0.174, 0.106], p=0.623; all other p>0.35). Likewise, participant motion during DWI acquisition had no effect on performance, measures of tract integrity, or the relationship between CA_3_-CA_1_ integrity and exception learning (see Methods, **Linking learning behaviour and neural measures**). None of the tract integrity, volume, or motion measures were significantly related to performance for similar items (all p>0.078). Additionally, exploratory analyses of diffusion tensor metrics (FA, MD, AD, and RD) and apparent fibre density (AFD) were not related to exception learning for either pathway (all p>0.19; see Supplemental Analysis).

## Discussion

Our results are consistent with a specific link between TSP integrity (characterized as CA_3_-CA_1_ white matter connections) and categorization performance for exception items. To our knowledge, this is the first empirical demonstration of the functional significance of CA_3_-CA_1_ integrity in the human brain. That is not to say, however, that our result is surprising; to the contrary, they are grounded in—and converge nicely with—simulations from computational models (Schapiro et al., 2017) that highlight a role for TSP in representing distinct episodes (but failing to capture cross-event regularities). Importantly, such representations were found across multiple paradigms—statistical learning, associative inference, and community structure learning—suggesting the generality of this function. Here, we build upon these previous results to ask whether TSP tract integrity is associated with specific aspects of performance in a different task: namely, category learning (Zeithamova et al., 2019). We set up our category structure such that a minority of items (“exceptions”) necessitated the use of specific memories for successful categorization, because they did not follow the broader rule. We also theorized that exceptions would benefit most from mismatch-driven encoding processes—a computation that requires CA_3_-based pattern completion—as they deviate most from memory-based expectations (Neunuebel and Knierim, 2014). Consistent with our hypotheses, we found that greater tract integrity for CA_3_-CA_1_ (but not ERC-CA_1_) was associated with superior performance for exceptions (but not the rule-following, similar items).

We did not find evidence for an association between MSP tract integrity (operationalized as ERC-CA_1_) and behaviour on either similar or exception items. Given that performance on the similar items might benefit from extraction of regularities across experiences—an operation purportedly supported by MSP (Schapiro et al., 2017)—we might have expected stronger MSP connections to be associated with better performance on *similar* items. It might be the case that certain features of our task reduced our chances of observing such an effect. For example, more repetitions (given the slower learning rate of CA_1_) or a different stimulus structure (e.g., in which there exist more “similar” items that accentuate the clustering around the anchors) might uncover evidence of such a relationship. These predictions remain to be tested in future work. It is also important to note that our characterization of MSP relies on ERC, a region susceptible to issues of signal-to-noise and decreased contrast with MRI methods. It is possible that our streamline estimation of ERC-CA_1_ was impacted by these imaging limitations to a greater extent than was CA_3_-CA_1_. Although the current findings point to a robust relationship between CA_3_-CA_1_ streamlines and exception learning, future work with tailored learning paradigms and imaging protocols optimized for ERC may provide a better characterization of MSP’s relationship to category learning.

Methodologically speaking, our study demonstrates the value of measuring the connectivity between hippocampal subregions in healthy human participants. There is limited work measuring structural properties of connections within the human hippocampus (Auhustinack, 2010; Yassa et al., 2010, 2011; Zeineh et al., 2012, 2017) and most of it has been done *post mortem* (Auhustinack, 2010; Zeineh et al., 2017). While informative from a basic science perspective, to fully test the predictions of computational models—which highlight the unique behavioural contributions of each pathway—we must study living human participants and carefully measure their behaviour on a theoretically informed choice of tasks. Previous reports interrogating the structure of the perforant path (part of TSP) in healthy aging have quantified fractional anisotropy (FA)—a diffusion-based metric describing the degree to which diffusion is directional in a particular region—within a combined dentate gyrus and CA_3_ region (Yassa et al., 2010, 2011). FA in this region was related to behavioural evidence for pattern separation, consistent with hypotheses (Yassa et al., 2011); however, this was observed in a very small group of older adult participants (N=11) due to the intensive data collection required and used a general metric of integrity that did not capture subfield-to-subfield connections specifically. We suggest that our approach using a smaller number of high-resolution DWI scans in healthy young adults improves feasibility and therefore opens the door to future, larger scale investigations testing predictions about which specific aspects of memory are related to anatomical properties of TSP-related connections. In addition, using such an approach in neuropsychological studies with clinical populations in the future may yield even greater variability and prove a useful and sensitive tool in understanding the neural basis of memory impairments specifically, and perhaps deficits in flexible cognition more generally.

Our work adds to a growing body of evidence suggesting that individual differences like intrinsic (i.e., task-free) functional connectivity (Elliott et al., 2019; Finn et al., 2020) and structural properties (Llera et al., 2019) relate to cognition in meaningful ways. Our results join prior research in highlighting the greater sensitivity of connectivity relative to volume-based measures (Yassa et al., 2010; Zajac et al., 2020). Here, we took a hypothesis-rather than data-driven approach to asking these questions; our analysis was limited to just two hippocampal pathways of greatest expected significance: CA_3_-CA_1_ as related to TSP, and ERC-CA_1_ as related to MSP. Moreover, we demonstrate these relationships outside the domain of a traditional episodic memory task. As mentioned earlier in the paper, our framework suggests that encoding of exceptions in a category learning context may—to an even greater degree than more standard one-shot learning tasks—benefit from a mechanism in which existing knowledge is reinstated and compared with current input (Love et al., 2004; Sakamoto and Love, 2004; Davis et al., 2012; Schapiro et al., 2017). It is therefore possible that performance on this category learning task may be *more* tightly linked to TSP-related function than would be a traditional episodic memory test that does not require such fine-grained discrimination among similar exemplars and a differentiation of related memories. This speculation remains to be empirically tested by future work.

Important limitations to the current findings are those inherent to using DWI to estimate white matter structure. Although state-of-the-art methods for segmenting MTL subregions (Yushkevich et al., 2015) and estimating white matter response and tractography (Smith et al., 2012, 2013; Tournier et al., 2019) were utilized to best approximate the underlying white matter structure, there remains the key caveat that in vivo DWI cannot discriminate TSP and MSP at the level of precision possible with ex vivo imaging and histology. The MTL is a crowded neural neighbourhood with interwoven grey and white matter, crossing white matter fibres, and surrounding ventricles. Given these challenges, performing tractography in MTL requires an approach that considers tissue types (Smith et al., 2012), incorporates crossing fibres in its underlying estimation (Tournier et al., 2012), and filters resulting streamlines according to biological plausibility (Smith et al., 2013, 2015). We incorporated all these key steps in the current work. Additionally, we acquired diffusion data with isotropic voxels (Zeineh et al., 2012) to avoid biases in tractography estimation (Jones and Leemans, 2010; Jones et al., 2013), corrected susceptibility distortions with reverse-phase encoded b_0_ structural images (Andersson et al., 2003; Smith et al., 2004), and collected two separate diffusion volumes for each participant to assess generalizability across acquisitions. One drawback to our approach is that the resolution of our diffusion acquisition was 2mm. We opted for a more standard resolution to increase SNR and provide whole brain coverage for tractography estimation; however, a higher resolution sequence coupled with multiple acquisitions (Zeineh et al., 2012) would likely provide better estimates of diffusion signal from distinct hippocampal pathways. Also, as is true of tractography in general, estimated streamlines are at best an approximation of the underlying distribution of white matter fibres and do not represent actual anatomy. Future work incorporating higher resolution diffusion (Yassa et al., 2010; Zeineh et al., 2012) and structural imaging along with advances in multi-shell diffusion acquisition (e.g., Pines et al., 2020) and neurite orientation dispersion and density imaging (NODDI) measures of microstructure (Zhang et al., 2012) will further bridge the divide between white matter anatomy and its estimation with MRI.

Our field is moving towards more fully describing the myriad ways in which the hippocampus— a structure once ascribed a relatively narrow, memory-only function—shapes cognition, broadly speaking (Shohamy and Turk-Browne, 2013). We suggest that future researchers considering its functional or structural connectivity might focus not only on how the hippocampus interacts with neocortical structures, but also how its subfields are interconnected with one another (Schapiro et al., 2017). Indeed, our findings underscore this internal circuitry of the hippocampus—rather than its sheer size or even subfield makeup—as the measurable anatomical property most tightly coupled with its function.

## Methods

### Participants

Forty-three volunteers (24 females, mean age 23.5 years old, ranging from 19 to 33 years) participated in the experiment. All subjects were right-handed, had normal or corrected-to-normal vision, and were compensated $20/hour for participating. Data from six participants were excluded from analysis due to technical issues with MRI acquisition (N=2) or experiment presentation (N=1), failure to follow instructions for the category learning task (N=2), and inability to estimate white matter streamlines (N=1).

### Category learning task

After an initial screening and consent in accordance with our University of Toronto Research Ethics Board-approved protocol, participants were instructed on the category learning task. Participants then performed the task while laying supine in the MRI scanner and viewing visual stimuli back-projected onto a screen through a mirror attached onto the head coil. Foam pads were used to minimize head motion. Stimulus presentation and timing was performed using custom scripts written in Matlab (Mathworks) and Psychtoolbox (www.psychtoolbox.org).

Participants were instructed to use feedback displayed on the screen to learn to classify visual images of cartoon flowers (**Fig. 1**) as growing better in the sun or the shade by considering its features. There were three feature dimensions: outer petal shape, inner petal shape, and colour/texture of flower centre, each of which could take on two values. Once in the MRI scanner, participants were reminded of the task instructions and told which of two buttons to press for each flower category.

To perform optimally in the learning task, participants had to pay attention to the combination of all three feature dimensions. Class associations were defined according to a rule-plus-exception category structure (Shepard et al., 1961): each category included an “anchor”, two “similar” items that differed from the anchor in only one feature, and one “exception” which differed from its category anchor in two features. However, exceptions differed from the alternate category anchor in only one feature, meaning they were in fact more visually similar to the anchor from the other flower category than to their own category anchor (see **Table 1**).

**Table 1:** Stimulus features, category association, and type for the category learning task. Each of the eight stimuli were represented by the binary values of three feature attributes. Based on their feature values, stimuli were category anchors, similar to their category anchor, or exception items.

| stimulus | feature dimension |  |  | category | type |
| --- | --- | --- | --- | --- | --- |
|  | 1 | 2 | 3 |  |  |
| 1 | 0 | 0 | 0 | A | anchor |
| 2 | 0 | 0 | 1 | A | similar |
| 3 | 0 | 1 | 0 | A | similar |
| 4 | 0 | 1 | 1 | B | similar |
| 5 | 1 | 0 | 0 | B | exception |
| 6 | 1 | 0 | 1 | A | exception |
| 7 | 1 | 1 | 0 | B | similar |
| 8 | 1 | 1 | 1 | B | anchor |

On each trial of the learning task (**Figure 1**), participants were shown a flower stimulus for 3.5s, during which time they were instructed to respond indicating the flower’s category by pressing one of two buttons on an MRI-compatible response box. Flower images subtended 7.3° × 7.3° of visual space. The stimulus presentation period was followed by a 0.5-4.5s fixation. A feedback screen consisting of the flower image, text of whether the response was correct or incorrect, and the correct category was shown for 2s followed by a 4-8s fixation. The frequency of the different stimulus types was purposefully unbalanced to make similar and exception stimuli rare events during learning. This was done to increase the difficulty of the task and to place a greater demand on generalizing learning of the anchors to similar and exception items. Within one trial block, each anchor was presented six times, the four similar items each four times, and the two exceptions each twice for a total of 24 trials. Trial order was randomly selected for each trial block. Participants completed six trial blocks for a total of 144 trials. The entire category learning task lasted approximately 40 minutes. Functional MRI data was collected during the learning task, but this data is not considered in the current study.

### Learning behaviour analysis

Participant-specific learning behaviour was characterized by the proportion of correctly categorized trials in the last two trial blocks separately for anchors, similar items, and exceptions (**Fig. 1**). The last two blocks were chosen such that a minimum of eight trials were used for accuracy calculations across the stimulus types in order to provide stable estimates of performance, as has been done previously (Nosofsky et al., 1994; e.g., Davis et al., 2012; Mack et al., 2020). Learning performance was evaluated with a mixed-effects linear regression that included a fixed effect of stimulus type and random intercepts for participants (*R* version 4.0.0, *lme4* version 1.1-23).

### MRI acquisition

Whole-brain imaging data were acquired on a 3.0T Siemens Prisma system at the Toronto Neuroimaging Facility housed at the University of Toronto. A high-resolution T1-weighted MPRAGE structural volume (TR = 2s, TE = 2.4ms, flip angle = 9°, FOV = 256mm, matrix = 256×256, 1mm iso-voxels) was acquired for co-registration and parcellation. Two oblique coronal T2-weighted structural images were acquired perpendicular to the main axis of the hippocampus (TR = 4s, TE = 66ms, matrix = 512×512, 0.43×0.43mm in-plane resolution, 2mm thru-plane resolution, 40 slices, no gap). Functional images were also acquired using a T2*-weighted EPI pulse sequence during the category learning task and during resting-state scans prior to and after the learning task. This functional data is not considered in the current study. Diffusion weighted images (DWI) were also collected in two separate scans one before and one after the learning task (axial echo-planar imaging, GRAPPA=2, multiband factor=2, b-value=1000, TR=4000ms, TE=70ms, 64 directions, 22cm FOV, 2mm-iso voxels, 70 slices, acquisition time=298s; sequence also included 4 repetitions of b-value=0 images). An additional two reverse-phase encoded b-value=0 images were collected for distortion correction.

### Regions of interest

Hippocampal subfields and medial temporal cortex subregions were automatically labelled using Automated Segmentation of Hippocampal Subfields (Yushkevich et al., 2015) and the Princeton Young Adult 3T ASHS Atlas (Hindy et al., 2016). Outputs from this procedure provided participant-specific regions of interests that included hippocampal subfields DG, CA_1_, a combined CA_2_ and CA_3_ regions (labelled here as CA_3_), and Subiculum and MTL cortical regions of ERC, PRC, and PHC separately for left and right hemispheres. Because here we focus only on the intra-hippocampal pathways and considered MTL cortex only in order to define TSP and MSP, we used only ERC (i.e., not PRC or PHC) in our analysis as the input structure to hippocampus and a component of MSP. These ROIs were co-registered to participants’ T1 anatomical space using the linear transformations provided by ASHS.

### Regional volume analysis

The volume of each hippocampal subfield and ERC was calculated from the ASHS-defined ROIs. To correct for overall head size differences across participants, intracranial volume (ICV) was estimated using the standard recon-all protocol in Freesurfer (version 6.0; Fischl et al., 2002) and entered as a predictor of ROI volume in separate linear regression models for each ROI (R version 4.0.0). The residuals of these models served as an ICV-corrected index of ROI volume (Mathalon et al., 1993) and were used in all analyses linking volume to learning performance as described below.

### Diffusion weighted imaging and tractography analysis

Our approach to characterizing white matter connectivity of intrinsic hippocampal pathways was to 1) estimate tractography at the whole-brain level, 2) use the whole-brain tractography to construct a structural connectome among the MTL ROIs described above, and 3) from the MTL-based connectome characterize the integrity of MSP and TSP with a relative measure of streamline counts. By starting with an estimation of whole brain tractography, our aim was to capture the global structure of white matter in order to reduce potential artificial constraints imposed by a limited field of view or misclassification of long-range fibres that pass through our targeted region but nonetheless have endpoints outside of MTL. This approach, the details of which are explained below, was applied separately to each of the two DWI scans collected from each participant.

Our approach departs from traditional DWI analyses that rely on diffusion tensor metrics of diffusivity (e.g., fractional anisotropy, mean diffusivity). Such metrics rely on estimates of diffusion tensors which are poor models of complex local fibres: they cannot represent more than one independent orientation nor account for crossing fibres (Tournier et al., 2012; Jeurissen et al., 2013; Jones et al., 2013; Riffert et al., 2014). Recent methodological advances have overcome these challenges through a constrained spherical deconvolution method that estimates the distribution of white matter fibre orientations within each voxel (Tournier et al., 2004, 2007, 2012, 2019). These distributions of fibre orientation can then be leveraged to estimate streamline tractography that better characterizes more complex fibre arrangements. The result is an estimate of white matter connectivity that more closely represents the underlying anatomical structure than traditional diffusion tensor metrics (Smith et al., 2013, 2015; Jeurissen et al., 2014b; Roine et al., 2015). As such, a constrained spherical deconvolution method was utilized in the current DWI analyses.

All DWI analyses were conducted with MRtrix (version 3.0 RC3) (Tournier et al., 2019) and FSL tools. First, for each participant, the average of the b_0_ volumes collected during DWI acquisitions were co-registered to the T1 anatomical volume with boundary based registration (Greve and Fischl, 2009) in FSL flirt using the white matter probability volume calculated in fmriprep (Esteban et al., 2019). This registration was used to register the diffusion data to the participant’s T1 anatomical volume. Diffusion data was denoised (Veraart et al., 2016) and pre-processed and corrected for distortions with FSL EDDY (Smith et al., 2004; Andersson and Sotiropoulos, 2016). Scan-specific estimates of motion during DWI acquisition (two per participant) were calculated from the EDDY motion correction outputs (Taylor et al., 2016). Specifically, the average of the volume-by-volume root-mean-squared errors (RMSE) across all voxels was calculated separately for each of the DWI scans. These motion indices were included in all analyses linking white matter tracts to learning behaviour as described below. The multi-tissue constrained spherical deconvolution (MSMT-CSD) framework was used to estimate the fiber orientation distribution (FOD) in each voxel (Jeurissen et al., 2014a). This framework estimates orientation distribution functions for different tissue types to create a signal distribution map of white matter, grey matter, and cerebrospinal fluid within the diffusion images. Whole-brain probabilistic tractography was estimated using the iFOD2 algorithm with 10 million streamlines, curvature threshold of 45 degrees, minimum fiber length of 10 mm, maximum fiber length of 250 mm, and seeds from the gray matter-white matter interface (Smith et al., 2012). Spherical-deconvolution informed filtering (SIFT) was then applied to match the streamline densities with the FOD lobe integrals (Smith et al., 2013). This step removed likely false positive tracts and reduced the number of streamlines to 2 million to create more biologically meaningful estimates of white matter fibres (Smith et al., 2013).

Participant-specific connectomes based on streamlines that connected the MTL and hippocampal ROIs were generated with MRtrix *tck2connectome*. To correct for individual differences in participants’ overall streamline counts, we scaled streamline counts to the proportion of all streamlines connecting MTL ROIs for each participant. This streamline count proportion, therefore, takes into account potential differences in streamlines due to ROI size and overall streamline density in order to target the relative comparison of TSP- and MSP-related WM streamline connections.

Whereas MSP involves a direct connection between ERC and CA_1_, the TSP circuit involves a multi-step path linking ERC to DG and CA_3_ before reaching CA_1_. Thus, to approximate connections along MSP and TSP with similar constraints (e.g., each with one start and endpoint) and put the two pathways on more equal footing in the analysis, also selecting them to be roughly the same length given the known distance-dependent motion artifacts in tractography (Baum et al., 2018), we focused on just the connections of CA_1_ with ERC and CA_3_ as indices of overall MSP- and TSP-related path integrity, respectively. Specifically, MSP-related path integrity was defined as the streamline count proportion connecting ERC and CA_1_; for TSP-related paths, it was the streamline proportion between CA_3_ and CA_1_. Output connections via fornix were not included in either pathway definition. It is important to note that at the resolution of DWI, diffusion signal from the entire extent of MSP and TSP would spatially overlap. Our choice to focus on distinct inputs to CA_1_ originating from different regions approximates MSP- and TSP-related connections at points where they are spatially separable. Note also, this definition of TSP-related connections is consistent with computational studies that characterized distinct functional properties of hippocampal pathways by down-weighting the connection between CA_3_ and CA_1_ in order to simulate a lesioned TSP (Ketz et al., 2013; Schapiro et al., 2017). Raw streamline counts for the two pathways (**Fig. 2B**) were not correlated across participants (r=0.167, p=0.323). To provide a comparison, we also defined streamline counts for uncinate fasciculus (UF). Specifically, we followed the same methods as described for the HPC pathways but estimated the number of streamlines connecting key UF seed regions of a combined lateral and medial orbitofrontal cortex and amygdala (**Table 2**). These regions were defined from the Desikan-Killiany atlas parcellation as part of participant-specific freesurfer outputs.

**Table 2:** Summary of streamline counts from HPC-related pathways and a comparison tract, uncinate fasciculus. Summary measures include the range, mean, and standard deviation of streamline counts across participants.

|  | range | mean (SD) |
| --- | --- | --- |
| CA<sub>3</sub>-CA<sub>1</sub> (TSP-related) | 44-382 | 191.1 (63.5) |
| ERC-CA<sub>1</sub> (MSP-related) | 3-178 | 70.4 (57.9) |
| uncinate fasciculus | 63-494 | 240.1 (99.1) |

### Quality control

The streamline estimation pipeline was evaluated for quality control at three different points. First, the hippocampal subfield and MTL cortical segmentations provided by ASHS were visually inspected for each participant. Participant-specific volume masks for the relevant ROIs were displayed on the high resolution T2 and then the T1 anatomical volumes to ensure valid segmentation. Second, the DWI-T1 registration results were visually inspected by transforming the average b0 volume from the DWI acquisition to T1 space and displaying the transformed volume along with the T1 volume. Although we checked for whole brain alignment, we specifically focused on the MTL region to ensure proper registration between the two imaging modalities. Third, for each participant, the streamline estimations from MRtrix were visually inspected by viewing the TSP-related (CA_3_-CA_1_) and MSP-related (ERC-CA_1_) streamlines in both 2D and 3D along with the ASHS-defined ROI masks and anatomical volumes. We ensured that streamlines had appropriate endpoints in the relevant ROIs and that the direction and path of the streamlines were consistent with the known anatomical properties of the white matter connections between CA_1_, CA_3_, and ERC. One participant was excluded due to failed streamline estimation. Authors M.G. and M.L.M. performed all visual inspections.

### Linking learning behaviour and neural measures

We assessed the relationship between learning behaviour and neural measures of CA_3_-CA_1_ and ERC-CA_1_ integrity and ROI volume with linear regression modelling (*R* version 4.0.0; *stats::lm* version 4.0.0; *lme4* version 1.1-23). To control for potential confounds of participant-specific motion during the DWI scans (Taylor et al., 2016) on our pathway integrity measures, a source of structured variability that could reasonably relate to behavioural indices of learning, we included an estimate of DWI scan motion averaged across the DWI scans as a predictor in all regression models involving pathway integrity. We found that DWI motion 1) was not correlated with either CA_3_-CA_1_ or ERC-CA_1_ integrity, 2) was not correlated with learning performance, and 3) did not change any relationships between pathway integrity and learning. The DWI motion estimate was included in all reported regression models that follow.

For the analysis reported in the main text, we averaged measures of CA_3_-CA_1_ and ERC-CA_1_ integrity from the two DWI scans and across hemispheres. The resulting data provided single measures of CA_3_-CA_1_ and ERC-CA_1_ integrity for each participant. These pathway integrity measures were simultaneously related to learning performance for both exceptions and similar items in separate linear models (see Results section). Our choice to average across DWI scans and hemispheres was motivated by the outcome of linear mixed effects model relating CA_3_-CA_1_ and ERC-CA_1_ integrity to exception learning performance that, in addition to DWI motion, included separate fixed effects factors for scan and hemisphere. The result of this model showed no effects for either factor (scan: β=-0.005, CI_95%_=[−0.033, −0.022], p=0.720; hemisphere: β=− 0.001, CI_95%_=[−0.029, −0.029], p=0.979). Importantly, while the main results show brain-behaviour relationships when collapsing across run and hemisphere, the significant relationship between CA_3_-CA_1_ integrity and exception learning performance was also observed when considering data from each DWI scan (repetition 1: β=0.123, CI_95%_=[0.038, 0.208], p=0.006; repetition 2: β=0.077, CI_95%_=[0.001, 0.155], p=0.050) and hemisphere (left: β=0.122, CI_95%_=[0.021, 0.223], p=0.020; right: β=0.073, CI_95%_=[0.004, 0.141], p=0.038) independently.

To evaluate the reliability of the main findings relating exception learning to CA_3_-CA_1_ and ERC-CA_1_ integrity, we conducted a 1) bootstrap resampling procedure and 2) robust regression analysis. For the bootstrap resampling analysis, on each iteration, data from participants was randomly sampled with replacement, the regression model evaluated, and the coefficients for CA_3_-CA_1_ and ERC-CA_1_ were saved. This procedure was repeated for 10,000 iterations to generate a distribution of coefficients for the effects linking pathway integrity to exception learning. The resulting coefficient distributions were used to calculate 95% confidence intervals and a p_boot_ statistic (**Fig. 2B**) that characterizes the proportion of resampling tests that resulted in a coefficient with a sign opposite of the distribution’s median. For robust regression, we estimated a robust regression model relating exception learning to CA_3_-CA_1_ and ERC-CA_1_ integrity and DWI motion estimates (*MASS::rlm* version 7.3-51.6, M-estimator with median-absolute deviance estimation). Importantly, the TSP effect remained (β=0.118, CI_95%_=[0.043, 0.193], p=0.004). Findings from both of these analyses suggest that the CA_3_-CA_1_ effect on exception learning is reliable and not driven by the contributions of outliers.

#### Direct comparison of CA_3_-CA_1_ and ERC-CA_1_ effects

An additional model was estimated that considered pathway integrity as the outcome variable with exception learning performance, pathway (CA_3_-CA_1_ vs. ERC-CA_1_), and their interaction as predictors. Importantly, the interaction term was significant (β=0.180, CI_95%_=[0.044, 0.317], p=0.010), suggesting that the relationship between CA_3_-CA_1_ integrity and performance was reliably stronger than the relationship between ERC-CA_1_ and performance. A similar model considering pathway integrity in light of learning performance for similar items showed no relationship for either pathway (all ps>0.123).

#### No relationship between volume and behaviour

The relationship between ROI volume and learning performance was evaluated with a linear regression model with ICV-corrected volumes for each hippocampal ROI (CA1, CA3, DG, SUB) and ERC entered as predictors of accuracy, separately for exceptions and similar items. Performance for neither item types was explained by individual differences in volume (all ps>0.16).

#### CA_3_-CA_1_ effects remain when controlling for volume

We also entered ROI volumes into a linear model along with CA_3_-CA_1_ and ERC-CA_1_ integrity and DWI scan motion as predictors of learning performance. In this extended model, the relationship between TSP integrity and exception learning remained reliable (β=0.129, CI_95%_=[0.021, 0.237], p=0.021) and no other predictors were significant (all ps>0.35). A model using the same predictors for similar item learning showed no effects (all ps>0.078).

#### CA_3_-CA_1_ effects remain when controlling for participant age and sex

We additionally entered participant sex and age as covariates along with CA_3_-CA_1_ and ERC-CA_1_ integrity and DWI scan motion as predictors of learning performance. Neither age (p=0.345) nor sex (p=0.369) were related to learning performance; additionally, the relationship between CA_3_-CA_1_ streamlines and exception learning remained (β=0.118, CI_95%_=[0.021, 0.214], p=0.018). There was also no evidence of simple linear relationships between age and sex and pathway integrity (all ps>0.2).

#### CA_3_-CA_1_ effects remain when including all TSP-related connections

We also considered TSP-related streamlines between all three steps in the pathway (ERC-DG-CA_3_-CA_1_) by summing the streamline counts between each pair of subregions. Consistent with the central findings, we found a significant relationship between streamline counts associated with this expanded definition of TSP-related white matter connections (β=0.102, CI_95%_=[0.019, 0.186], p=0.017).

#### No relationship between DWI scan motion and behaviour or pathway integrity

To fully consider the effect of motion during DWI scan collection, we assessed the simple linear relationship between motion and exception learning, and CA_3_-CA_1_ and ERC-CA_1_ integrity. Motion had no effect on any of these variables (all ps>0.45).

## Acknowledgements

Thanks to Chanel Fu and Anna Blumenthal for assistance with data collection; and to Dasa Zeithamova and members of the Mack and Budding Minds Labs for helpful discussions. This research was supported by Natural Sciences and Engineering Research Council (NSERC) Discovery Grants (RGPIN-2017-06753 to MLM and RGPIN-2018-04933 to MLS), Canada Foundation for Innovation and Ontario Research Fund (36601 to MLM and 36876 to MLS), and a Brain Canada Future Leaders in Canadian Brain Research Grant (MLM).

## Footnotes

1 The same positive relationship between TSP-related streamlines and exception learning was found when considering the sum of white matter connections across additional parts of TSP (i.e., ERC-DG, DG-CA_3_, CA_3_-CA_1_, β=0.102, CI_95%_=[0.019, 0.186], p=0.017).

